# TEMINET: A Co-Informative and Trustworthy Multi-Omics Integration Network for Diagnostic Prediction

**DOI:** 10.1101/2024.01.03.574118

**Authors:** Haoran Luo, Hong Liang, Hongwei Liu, Zhoujie Fan, Yanhui Wei, Xiaohui Yao, Shan Cong

## Abstract

Advancing the domain of biomedical investigation, integrated multi-omics data have shown exceptional performance in elucidating complex human diseases. However, as the variety of omics information expands, precisely perceiving the informativeness of intra- and inter-omics becomes challenging due to the intricate interrelations, thus posing significant obstacles in multi-omics data integration. To address this, we introduce a novel multi-omics integration approach, referred to as TEMINET. This approach enhances diagnostic prediction by leveraging an intra-omics co-informative representation method and a trustworthy learning strategy used to address inter-omics fusion. Considering the multifactorial nature of complex diseases, TEMINET utilizes intra-omics features to construct disease-specific networks, then applies graph attention networks and a multi-level framework to capture more collective informativeness than pairwise relations. To perceive the contribution of co-informative representations within intra-omics, we design a trustworthy learning strategy to identify the reliability of each omics in integration. To integrate inter-omics information, a combined beliefs fusion approach is deployed to harmonize the trustworthy representations of different omics types effectively. Our experiments across four different diseases using mRNA, methylation, and miRNA data demonstrate that TEMINET achieves advanced performance and robustness in classification tasks.

## 1. Introduction

In light of recent advancements in the acquisition of high-throughput omics data, multi-omics integration has emerged as a rapidly expanding research field, aiming to provide a more comprehensive understanding of the underlying biological processes and molecular mechanisms involved in complex diseases [1,2]. Compared to single-omics studies, integrating multiple types of omics data enables the capture of complementary information across various molecular layers, leading to a more holistic view of biological systems [3]. Traditional approaches often involve statistical tools, which may have limited capacity to fully capture the complex, non-linear relationships present in multi-omics datasets. In recent years, the application of artificial neural networks (ANN) to multi-omics studies has emerged as a promising avenue for addressing these limitations [4–6]. ANN approaches can learn intricate patterns within the data, enabling more accurate predictions and classifications, as well as the identification of previously unexplored relationships between molecular entities. These methods have shown great potential in various biomedical applications, such as disease prediction, patient stratification, and the discovery of novel biomarkers [7].

Despite significant achievements, the stability and biological explainability of many existing multi-omics integration approaches remain underdeveloped, primarily because of the insufficient exploration of both intra- and inter-omics interaction information. Existing studies, which are divided into early, intermediate, and late integration strategies, exhibit a tendency to utilize the fitting capabilities of ANNs for information integration [7–9]. Early integration methods, such as FCNNs [10–12] and autoencoders (AE) [13–15], perform integration through concatenating representations. These approaches treat intra-omics as an individual molecule and inter-omics as an individual modality. When facing modalities with different distributions and highly homogeneous, they are less likely to perform effectively. Intermediate fusion emphasizes the interactions within inter-omics information, rather than perceiving each omics type individually. For example, variational AEs (VAEs) are widely used to perform a fusion of homogeneous data types in a joint manner [16,17]. Despite the advantages of coordinated representation learning in intermediate fusion, it operates under the presumption of equal contribution from each modality, which is challenging for the fitting capabilities of ANNs, especially in cases of feature imbalance, severe missing modalities, or substantial noise interference. Considering the uncertainty of such situations, trustworthy learning methodologies have been adopted, transforming the operation from feature embedding to a decision-level process, thereby stabilizing and enhancing the outcomes of late integration strategies. Han et al. [18] introduce a dynamic fusion approach for multi-modal classification. This method employs true-class probability to asses the classification confidence across different omics and then adjusts through modality confidence weighting for integration. Wang et al. [19] propose Mogonet, which employs a View Correlation Discovery Network (VCDN) to integrate initial classifications instead of fusing features across modalities, thereby utilizing label-correlated information in the shared space to produce final classification labels. We observe that by perceiving and integrating the informativeness of each modality and inter-modality from distinct samples in trustworthy multimodal learning, practical applications can be significantly enhanced. However, traditional intra-modality information embedding may inadequately capture the full scope of informativeness, primarily due to the biological explainability of intra-omics. Such studies may overlook essential aspects of the underlying biological processes, potentially leading to a limited understanding of the complex molecular networks driving various biological systems[20]. Ultimately, this could result in challenges in effective integration.

Acknowledging the interrelations of molecular functions is essential in omics research, due to the multifactorial nature of complex diseases [21,22], rather than viewing each genetic factor as an independent entity. To address these challenges, there is a growing number of investigations directed toward the construction of pre-established disease networks by reconfiguring omics data into graph-based structures, reflecting a growing recognition of the importance of contextualized molecular interactions in understanding disease mechanisms. For instance, Ramirez et al. [23] employ a GCN approach for cancer classification, leveraging a framework of gene co-expression based on Spearman correlating analysis. Althubaiti et al. [24] utilize the DeepMOCCA framework, combining omics data with protein-protein interaction networks for improved cancer prognosis predictions. Furthermore, Xing et al. [25] adopt an approach involving a weighted correlating method to build prior knowledge graphs. This method is particularly advantageous for unraveling disease-specific complexities. Therefore, inspired by the effectiveness of network representations in omics studies, we are motivated to adopt a graph-based representation to enhance the informativeness of intra-omics. This approach is intuitive, easy to interpret, and has the potential to better preserve the inherent structure and capture functional interactions from omics data, resulting in improved disease prediction performance.

Based on the above observations, we propose a multi-omics integration framework, named TEMINET, for disease predictive diagnosis that leverages graph attention networks and an uncertainty-based trustworthy strategy. Specifically, we construct a disease-specific network for each omics data to represent large-scale, unstructured, and irregular data effectively. We apply hierarchical graph attention networks to capture co-informative intra-omics representations. Then, a trustworthy learning mechanism is employed to assess the reliability of informativeness by using uncertainty information. Combine beliefs fusion integrates informativeness and uncertainty into an inter-omics fusion embedding for executing subsequent tasks. We conduct extensive experiments to show the effectiveness and robustness of TEMINET for multi-omics prediction. Our results demonstrate that the combination of graph-based feature representation and uncertainty-based trustworthy learning integration surpasses the state-of-the-art models on four disease classification tasks. We further employ a global interpretation approach to identify important biomarkers and analyze their disease-related functional relevance.

## 2. Materials and Methods

### 2.1 Overview of TEMINET

The proposed method is illustrated in Figure 1. It begins with constructing a coexpression graph for the omics data of each subject via the WGCNA. The second step involves generating initial classification results for each omics data through a multi-level GAT process, utilizing three layers to extract intra-omics features from the initial GAT network *G*_0_, with further updating in GAT networks *G*_1_ and *G*_2_. Thirdly, the uncertainty of each initial distribution is parameterized using subjective logic. Based on the Dempster-Shafer theory, the integration of multi-omics evidence comes from the inference of overall uncertainty and classification probability. The whole inference process concludes with the final label prediction of each subject.

**Figure 1.**
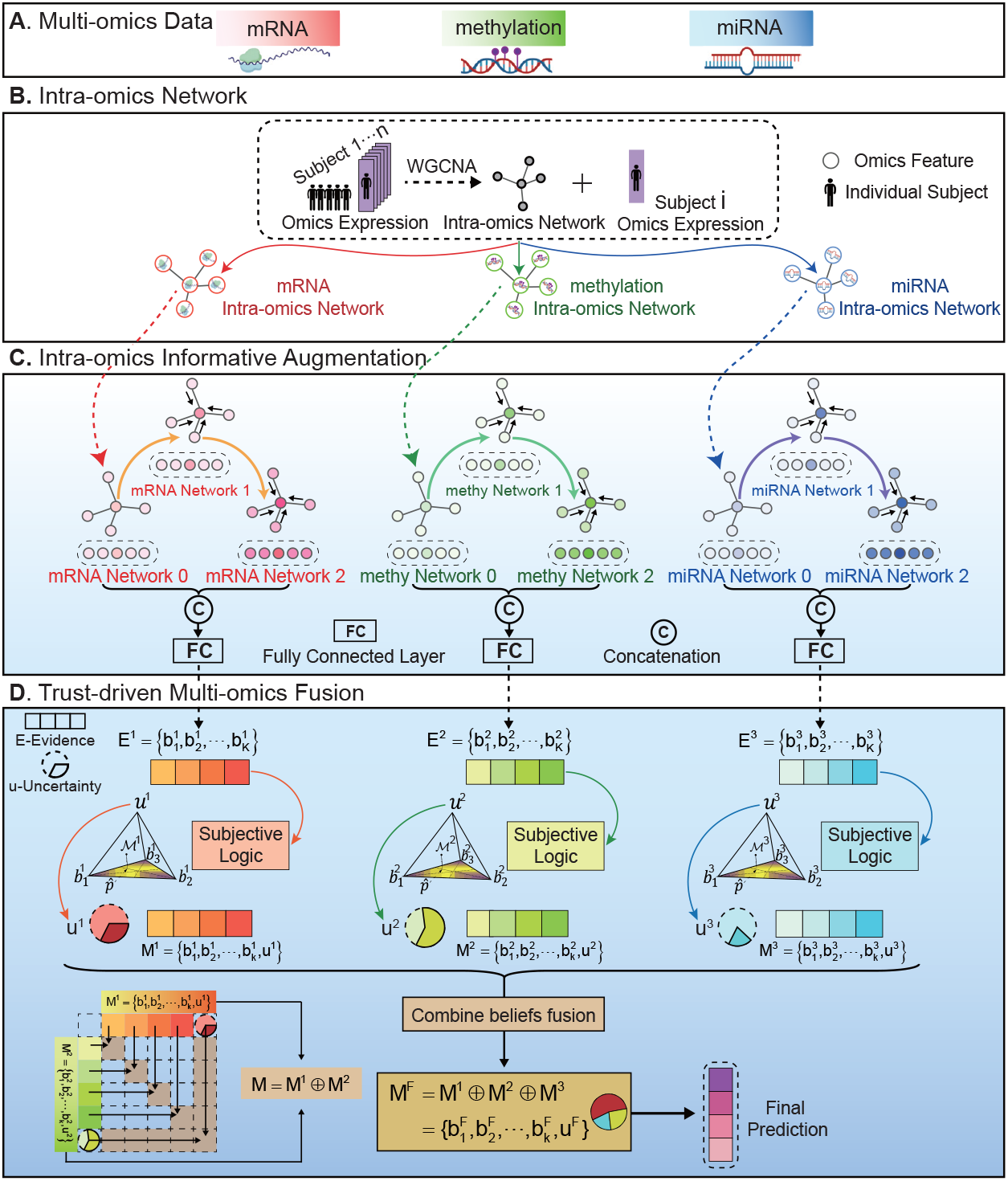
Framework of TEMINET. **(A)** TEMINET operates on a sample-wise basis with multi-omics information for each individual sample being imported into the model. **(B)** The first intra-omics network is built using the WGCNA. **(C)** The intra-omic information at each omics-level is augmented using the multi-level GAT. **(D)** The evidence is evaluated by the subject logic module to obtain uncertainty. During the integration phase, the trustworthy informativeness and uncertainty from each omics are amalgamated into a composite embedding encompassing inter-omics information. The fusing representation is subsequently applied to implement a downstream classification task.

### 2.2 Dataset Overview

In our investigation, we performed analysis using four public benchmark datasets provided by Wang et al. [19], including ROSMAP for binary classification (distinguishing between Alzheimer’s disease (AD) and normal control (NC)), BRCA for the classification of breast invasive carcinoma into PAM50 subtypes (including five categories), low-grade glioma (LGG) for distinguishing between grade *I I* and grade *I I I*, and KIPAN referring to the classification of renal cell carcinoma subtypes. The study utilized three categories of omics data: mRNA expression, DNA methylation, and miRNA expression. Table 1 provides comprehensive details regarding these datasets.

**Table 1.** Overview of datasets used in this investigation.

| Dataset | Source | Subjects | Number of origin data |  |  | Number of data for study |  |  |
| --- | --- | --- | --- | --- | --- | --- | --- | --- |
|  |  |  | mRNA | methy | miRNA | mRNA | methy | miRNA |
| ROSMAP | AMPAD | Normal Control: 169, Alzheimer’s Disease: 182 | 55,889 | 23,788 | 309 | 200 | 200 | 200 |
| BRCA | TCGA | Luminal A: 436, Luminal B: 147, HER2-enriched: 46, Normal-like: 115, Basal-like: 131 | 20,531 | 20,106 | 503 | 1,000 | 1,000 | 503 |
| LGG | TCGA | Grade II: 246, Grade III: 264 | 20,531 | 20,114 | 548 | 2,000 | 2,000 | 548 |
| KIPAN | TCGA | KICH: 66, KIRC: 318, KIRP: 274 | 20,531 | 20,111 | 445 | 2,000 | 2,000 | 445 |

### 2.3 Intra-omics Network Construction

The development of functional interaction networks is integral to understanding the pathogenesis of complex diseases. To leverage synergistic relationships among intra-omics molecules, the initial step in our methodology involved implementing Weighted Correlation Network Analysis (WGCNA [26]) to construct intra-omics co-expression networks, as shown in Figure 1(B). The construction of the intra-omics graph *G*_0_ for each subject involved several key stages: Firstly, an adjacency matrix was formed through WGCNA, with each entry indicating the correlation strength between pairs of omics features. Secondly, this matrix was transformed into an edge matrix by applying a threshold. Thirdly, a co-expression network was constructed for individual subjects, assigning omics data expression values to nodes as their features.

In this context, an initial co-expression graph network of each patient is denoted as 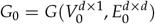. Here, *V*^*d×*1^ symbolizes the attributes of *d* nodes. *E*^*d×d*^ represents the edge matrix derived from the co-expression correlations computed via WGCNA. Specifically, for each subject *n*, a feature vector of dimension 1 *× d* is generated, where *d* represents the number of features. For a group of *N* subjects within a single omics data, an *N × d* matrix is formulated to calculate the co-expression matrix *A*^*d×d*^. The correlation *A*_*ij*_ between node *v*_*i*_ and *v*_*j*_ is given by

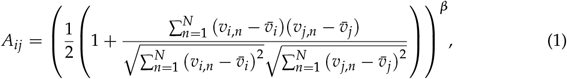

where 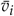 and 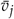denote the mean values of nodes *v*_*i*_ and *v*_*j*_, and *β* denotes an adjustable parameter set by WGCNA. The edge matrix 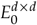 is then obtained by binarizing the values from the matrix *A*^*d×d*^. In conclusion, the construction of intra-omics co-expression networks is implemented for each dataset of mRNA, methylation, and miRNA, respectively.

### 2.4. Intra-omics Informative Augmentation

To leverage the information embedding in nodes and their associations within co-expression matrices effectively, we introduced the GAT [27] to enhance the disease-specific characteristics of the omics dataset. GAT incorporated masked self-attention based layers to enable dynamic weighting of neighboring node contributions, which allowed GAT to selectively focus on more pertinent adjacent nodes, thereby diminishing the impact of nodes that are lesser-significant. As a result, GAT exhibited a superior ability to discern intricate and non-structure connections, as well as variations within the topology of the graph.

Specifically, an initial network for each omics subject *n*, denoted as 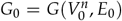 from Stage (B), is updated through a GAT layer. Initially, for a node *h*_*i*_ within this network, the normalization of attention coefficients *α*_*ij*_ with its neighboring nodes *h*_*j*_ are calculated as follows:

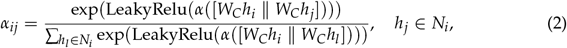

where ∥ is the concatenation operation, and *W*_*C*_ is the shared parameter matrix for linear transformation. To ensure a more stable self-attention update, a multi-head approach is introduced [27]. We conducted a process in which the attention-layer functions were implemented *T* times, each with unique parameters. The outcomes from these replications, indicated as *h*^*′i*^, are then conducted an aggregating concatenation in sequence as follows:

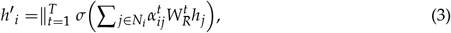

where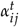 represents the attention coefficients from the *t*-th attention head, and *W*^*t*^ is the weight matrix associated with the *t*-th head.

To enhance the exploration of internal feature relationships, we have incorporated the multi-level feature representation approach implemented by Xing et al. [25]. The foundational network *G*_0_ encapsulates data corresponding to the primal features. Subsequently, *G*_1_ evolves from *G*_0_ via the deployment of a multi-head GAT attention-layer. Analogously, *G*_2_ is generated from *G*_1_. This progression through three progressive graph network layers creates a hierarchical integration structure, systematically amalgamating features across the GAT layers. Ultimately, the vectors produced from these transformation stages are fused, resulting in comprehensive feature representations. This design facilitates a dynamic optimization of feature interplays within the network, leading to a more substantial and comprehensive representation of the fundamental biology mechanisms. Using a similar process, we also implemented the DNA methylation and miRNA informative augmentation for each subject.

### 2.5 Trust-driven Multi-omics Fusion

In traditional multi-omics integration methods, the trustworthiness of different datasets is often not adequately considered, leading to potential inaccuracies in understanding complex biological processes. To address this, we introduced an uncertainty-based trustworthy learning approach in our integration method [28]. This approach is designed to enhance trustworthiness and precision in multi-omics data integration by quantifying inherent uncertainty in each modality. It leverages this measure of uncertainty to jointly perceive informativeness across intra- and inter-omics. Given that uncertainty assessments define confidence in predictions, the evidence of a dataset with lower uncertainty should obtain higher trustworthiness and be assigned a larger contribution to multi-omics integration.

Evidence in a classifier is generally considered the outputs of a neural network processed through an activation function like softmax. In our study, the evidence 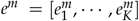 for the *m*^*th*^ omics category across *K* classes is generated from GAT-enhanced features. In the augmentation module, GAT-enhanced features are transformed into evidence through a sequence of fully connected layers and an active layer. Here, we set a cross-entropy loss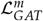 to modify the GAT augmentation module.

Secondly, to obtain the uncertainty, we applied subjective logic [29] to the evidence 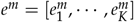. For each class *k* in the *m*^*th*^ omics category, the belief mass 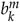and uncertainty mass *u*^*m*^ are calculated, ensuring 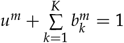. The concentration parameters *α*^*m*^ of the Dirichlet distribution are determined from the evidence, where 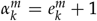. The belief mass for each class is computed as 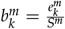and the overall uncertainty as 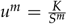, with 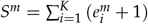. This formulation quantifies the confidence in each class prediction and the overall uncertainty in the classification for the *m*^*th*^ omics category. In summary, for the *m*^*th*^ omics category, the more evidence gathered for each of the *K* classes, the higher the probability assigned to the respective class, thus reducing uncertainty. Conversely, a scarcity of evidence leads to increased uncertainty. Utilizing subjective logic, this approach models second-order probabilities and uncertainties for the *m*^*th*^ omics category, effectively countering the overconfidence often seen in traditional neural network classifiers.

Thirdly, within the methodological framework for multi-omics fusion, we applied the Dempster-Shafer theory to synthesize evidence from different classes. This process consolidates independent probability mass assignments from each class into a unified joint mass. Dempster rule orchestrates this fusion to merge belief and uncertainty across the omics spectrum, symbolized as:

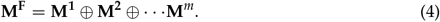

Taking the fusion of two omics categories as an example, where **M**^**1**^ is represented as 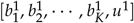and **M**^**2**^ as 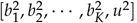, the fusion process is initiated as follows:

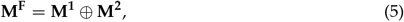

and

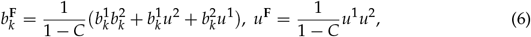

where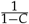 denotes the scale factor for normalization. After the 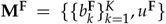 obtained, the final evidence can be inferred with *e*_*k*_ = *b*_*k*_ *× S* and 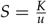.

Furthermore, we introduced an enhanced cross-entropy loss function by integrating sample-specific evidence as follows:

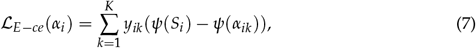

where *α*_*i*_ is the parameter of the Dirichlet distribution for the *i*th sample, and *ψ*(*·*) is the digamma function. Building on this, an overall sample-specific loss function, which combines the adjusted cross-entropy loss with a Kullback-Leibler divergence term to effectively manage the evidence for incorrect labels, is defined as:

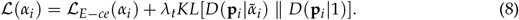

The 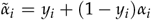is the modified attribute of the Dirichlet distribution, and *λ* is a balance factor greater than zero. This design helps penalize incorrect class evidence while preserving the evidence for the correct class. To ensure the informative augmentation and evidence fusion are updated simultaneously, a total loss is defined as:

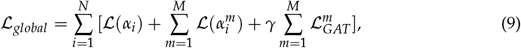

where *γ* denotes an adjusted attribute. We deployed *γ* = 1 across our investigations.

### 2.6. Identifying biomarkers with TEMINET

In the realm of biomedical research, the elucidation of biomarkers is pivotal for unraveling the intricacies of biological processes and providing insight into diverse outcomes. Concurrently, there is a growing need in clinical research for interpretable models that elucidate underlying disease mechanisms and bolster model credibility. Consequently, we applied a global interpretation method to identify the importance of biomarkers in our model. Specifically, to evaluate omics features in computational models, normalization of these features to a scale from zero to one is initially implemented. Feature ablation involves individually removing features to evaluate their impact on model efficacy, with a focus on classification capability. The importance of each feature is determined by observing the reduction in model performance post-ablation. In binary classification and multi-class classification tasks, the F1 score and F1 macro score are used to measure performance reduction resulting from feature ablation. This process is implemented with the best-performing model. For multi-omics data, we implemented this strategy in each type of omics data.

## 3. Results

Our research evaluates the effectiveness of the proposed model relative to current methodologies across diverse classification tasks. We also explore the advantages of incorporating additional omics datasets to enhance classification accuracy. In the comparative analysis of various methodologies, accuracy (ACC) is a common metric employed for binary and multi-class classifications. In addition to ACC, binary classifications also utilize the F1 score (F1) and the area under the receiver operating characteristic curve (AUC). For multi-class classifications, the ACC, the weighted average F1 score (F1_weighted), and the macro-averaged F1 score (F1_macro) are used. Our experimental framework replicates the same settings and provides the mean and standard deviation of these evaluative results.

Fourteen computational approaches are investigated, which include six classic classifiers employing early integration strategies: K-nearest neighbor classifier (KNN) [30], support vector machine (SVM) [31], Lasso [32], random forest (RF) [33], XGboost [34], and fully connected neural networks (NN) [35]. Furthermore, five multi-omics classifiers are analyzed: group-regularized ridge regression [36], BPLSDA for projecting data into latent structures with discriminant analysis [37], block PLSDA with additional sparse constraints (BSPLSDA) [37], concatenation of final representations (CF) [38] for late-stage multi-omics data, gated multimodal fusion (GMU) [39] that integrates intermediate representations, along with three advanced methods, Mogonet [19], MODILM [40], and Dynamic [18].

### 3.1. Classification performance comparison

In Table 2, a comparative analysis between the proposed model and established methods on ROSMAP and LGG datasets was provided. Compared to alternative approaches, the outcomes demonstrated that our suggested model performed better on binary classification tests. TEMINET outperformed the suboptimal method in many evaluation metrics, though not in the AUC metric for the ROSMAP dataset. The outcomes demonstrated the advantages of integrating informative data utilizing a multi-level graph-based attention framework and disease-specific networks. Moreover, the proposed model significantly exceeded the performance of MODILM (GAT). This advantage was likely due to the incorporation of uncertainty-based adaptive fusion, enhancing the capability of the model to select and utilize the informative modalities, thus ensuring a more precise characterization of each subject. As shown in Table 3, the proposed method continued to lead in overall performance, yet it showed a slightly lower efficacy in the five-class BRCA task when compared to the top-performing method in this specific area. This further indicated the strength of our model in leveraging disease-specific network analysis and graph attention mechanisms, underscoring its potential despite the room for improvement in certain multi-class classification tasks.

**Table 2.** Comparison with advanced methods on the ROSMAP and LGG datasets. Top-performing results are emphasized in boldface. Metrics marked with * signify a substantial improvement of our model over the suboptimal method, as evidenced by an independent t-test yielding *P <* 0.05.

| Method | ROSMAP (two categories) |  |  | LGG (two categories) |  |  |
| --- | --- | --- | --- | --- | --- | --- |
|  | ACC (%) | F1 (%) | AUC (%) | ACC (%) | F1 (%) | AUC (%) |
| KNN | 65.7±3.6 | 67.1±4.4 | 70.9±4.5 | 72.9±3.4 | 73.8±3.3 | 79.9±3.8 |
| SVM | 77.0±2.4 | 77.8±1.6 | 77.0±2.6 | 75.4±4.6 | 75.7±5.0 | 75.4±4.6 |
| Lasso | 69.4±3.7 | 73.0±3.3 | 77.0±3.5 | 76.1±1.8 | 76.7±2.2 | 82.3±2.7 |
| RF | 72.6±2.9 | 73.4±2.1 | 81.1±1.9 | 74.8±1.2 | 74.2±1.0 | 82.3±1.0 |
| XGBoost | 76.0±4.6 | 77.2±4.5 | 83.7±3.0 | 75.6±4.0 | 76.7±3.2 | 84.0±2.3 |
| NN | 75.5±2.1 | 76.4±2.1 | 82.7±2.5 | 73.7±2.3 | 74.8±2.4 | 81.0±3.7 |
| GRridge | 76.0±3.4 | 76.9±2.9 | 84.1±2.3 | 74.6±3.8 | 75.6±3.6 | 82.6±4.4 |
| BPLSDA | 74.2±2.4 | 75.5±2.3 | 83.0±2.5 | 75.9±2.5 | 73.8±3.1 | 82.5±2.3 |
| BSPLSDA | 75.3±3.3 | 76.4±3.5 | 83.8±2.1 | 68.5±2.7 | 66.2±3.0 | 73.0±2.6 |
| CF | 78.4±1.1 | 78.8±0.5 | 88.0±0.5 | 81.1±1.2 | 82.2±0.4 | 88.1±0.4 |
| GMU | 77.6±2.5 | 78.4±1.6 | 86.9±1.6 | 80.3±1.5 | 80.8±1.2 | 88.6±1.2 |
| Mogonet | 81.5±2.3 | 82.1±2.2 | 87.4±1.2 | 81.6±1.6 | 81.4±1.4 | 84.0±2.7 |
| MODILM | 84.3±1.2 | 85.0±0.8 | 89.1±1.2 | 82.8±0.7 | 82.5±2.3 | 86.1±1.5 |
| Dynamic | 84.2±1.3 | 84.6±0.7 | **91.2±0.7** | 83.3±1.0 | 83.7±0.4 | 88.5±0.4 |
| Ours | **87.9±0.4\*** | **88.2±0.4\*** | 87.6±1.3 | **84.3±0.7\*** | **84.1±0.9** | **88.6±1.2** |

**Table 3.** Comparison with advanced methods on the BRCA and KIPAN datasets. Top-performing results are emphasized in boldface. Metrics marked with * signify a substantial improvement of our model over the suboptimal method, as evidenced by an independent t-test yielding *P <* 0.05.

| Method | BRCA (five categories) |  |  | KIPAN (three categories) |  |  |
| --- | --- | --- | --- | --- | --- | --- |
|  | ACC (%) | F1-W<sup>1</sup> (%) | F1-M<sup>1</sup> (%) | ACC (%) | F1-W<sup>1</sup> (%) | F1-M<sup>1</sup> (%) |
| KNN | 74.2±2.4 | 73.0±2.3 | 68.2±2.5 | 96.7±1.1 | 96.7±1.1 | 96.0±1.4 |
| SVM | 72.9±1.8 | 70.2±1.5 | 64.0±1.7 | 99.5±0.3 | 99.5±0.3 | 99.4±0.4 |
| Lasso | 73.2±1.2 | 69.8±1.5 | 64.2±2.6 | 97.4±0.2 | 97.4±0.2 | 97.2±0.4 |
| RF | 75.4±0.9 | 73.3±1.0 | 64.9±1.3 | 98.1±0.6 | 98.1±0.6 | 97.5±1.1 |
| XGBoost | 78.1±0.8 | 76.4±1.0 | 70.1±1.7 | 99.3±0.8 | 99.3±0.8 | 98.9±1.4 |
| NN | 75.4±2.8 | 74.0±3.4 | 66.8±4.7 | 99.1±0.5 | 99.1±0.5 | 99.1±0.5 |
| GRridge | 74.5±1.6 | 72.6±1.9 | 65.6±2.5 | 99.4±0.4 | 99.4±0.4 | 99.3±0.4 |
| BPLSDA | 64.2±0.9 | 53.4±1.4 | 36.9±1.7 | 93.3±1.3 | 93.3±1.3 | 91.9±2.1 |
| BSPLSDA | 63.9±0.8 | 52.2±1.6 | 35.1±2.2 | 91.9±1.2 | 91.8±1.3 | 89.5±1.4 |
| CF | 81.5±0.8 | 81.5±0.9 | 77.1±0.9 | 99.2±0.5 | 99.2±0.5 | 98.8±0.9 |
| GMU | 80.0±3.9 | 79.8±5.8 | 74.6±5.8 | 97.7±1.6 | 97.6±1.7 | 95.8±3.2 |
| Mogonet | 82.9±1.8 | 82.5±1.6 | 77.4±1.7 | **99.9±0.2** | **99.9±0.2** | **99.9±0.2** |
| MODILM | 84.5±0.9 | 84.0±1.6 | 80.4±1.2 | 99.2±0.8 | 99.2±0.8 | 99.2±0.8 |
| Dynamic | 87.7±0.3 | **88.0±0.5** | 84.5±0.5 | **99.9±0.2** | **99.9±0.2** | **99.9±0.3** |
| Ours | **88.0±0.8** | 85.5±1.3 | **88.2±0.7\*** | **99.9±0.2** | **99.9±0.2** | **99.9±0.2** |

<sup>1</sup> The terms F1-M and F1-W denote the F1 macro and weighted scores, respectively.

### 3.2. Ablation comparison of model key component

In an extensive ablation study of our framework, as shown in Table 4, performance metrics were compared against established benchmarks. Our model incorporated an advanced uncertainty-based mechanism and omic-specific co-expression networks and achieved increases in ACC and F1, particularly within the ROSMAP dataset, where it outstripped the GAT+NN model by 4.3 % and 4.4%. However, there was a slight decrement in the AUC by 1.0% compared to the second-best models. This outcome suggested that while our model advanced predictive accuracy, it did so with a trade-off in AUC performance, signaling an area for further refinement in balancing predictive precision with generalizability across diverse datasets. The result of the LGG and KIPAN datasets also revealed improvements across all metrics, reflecting the exceptional ability of our model to capture intricate data patterns. For the BRCA dataset, our model exhibited enhancement in both ACC and the F1-M, yet it encountered a slight decrease in the F1-W score, indicating potential for further refinement in the equilibrium of precision and recall.

**Table 4.** The study examined key components of TEMINET on benchmark datasets, with the top-performing performances highlighted.

| Dataset | Metric | NN<sup>1</sup> | NN<sup>1</sup>+Trust | GAT+NN<sup>1</sup> | Our model |
| --- | --- | --- | --- | --- | --- |
| ROSMAP | ACC (%) | 76.6 ± 2.3 | 82.5±0.9 | 84.3±1.6 | **87.9±0.4** |
|  | F1 (%) | 77.7 ± 1.9 | 82.3±0.6 | 84.5±1.6 | **88.2±0.4** |
|  | AUC (%) | 81.9 ± 1.7 | **88.5±0.6** | **88.5±0.6** | 87.6±1.3 |
| LGG | ACC (%) | 74.0 ± 3.9 | 81.9±0.8 | 82.4±0.5 | **84.3±0.7** |
|  | F1 (%) | 75.6 ± 3.6 | 81.5±0.4 | 82.2±1.3 | **84.1±0.9** |
|  | AUC (%) | 82.4 ± 3.6 | 87.1±0.4 | 87.7±1.3 | **88.6±1.2** |
| BRCA | ACC (%) | 79.6 ± 1.2 | 84.2±0.5 | 87.7±0.4 | **88.0±0.8** |
|  | F1-W<sup>2</sup> (%) | 78.4 ± 1.4 | 84.4±0.9 | **88.2±0.5** | 85.5±1.3 |
|  | F1-M<sup>2</sup> (%) | 72.3 ± 1.8 | 80.6±0.9 | 84.8±1.7 | **88.2±0.7** |
| KIPAN | ACC (%) | 98.8 ± 1.1 | 99.7±0.3 | 98.9±0.4 | **99.9±0.2** |
|  | F1-W<sup>2</sup> (%) | 98.8 ± 1.1 | 99.7±0.3 | 98.9±0.5 | **99.9±0.2** |
|  | F1-M<sup>2</sup> (%) | 98.1 ± 1.6 | 99.4±0.5 | 98.7±0.7 | **99.9±0.2** |

<sup>1</sup> The NN refers to a neural network.
<sup>2</sup> The terms F1-M and F1-W denote the F1 macro and weighted scores, respectively.

### 3.3. Ablation study comparing integration performance across varied omics categories

In our investigation, we analyzed the effectiveness of different omics data combinations for classification performance. Figure 2 showed that using all three omics types outperformed combinations of just two. This result underlined the individual and substantial contributions provided by different omics categories. Moreover, it confirmed the advantage of integrating multiple omics approaches. Our results indicated that TEMINET enhanced classification performance by incorporating multi-omics informative data. Notably, these improvements were more pronounced with the progressive addition of various omics types.

**Figure 2.**
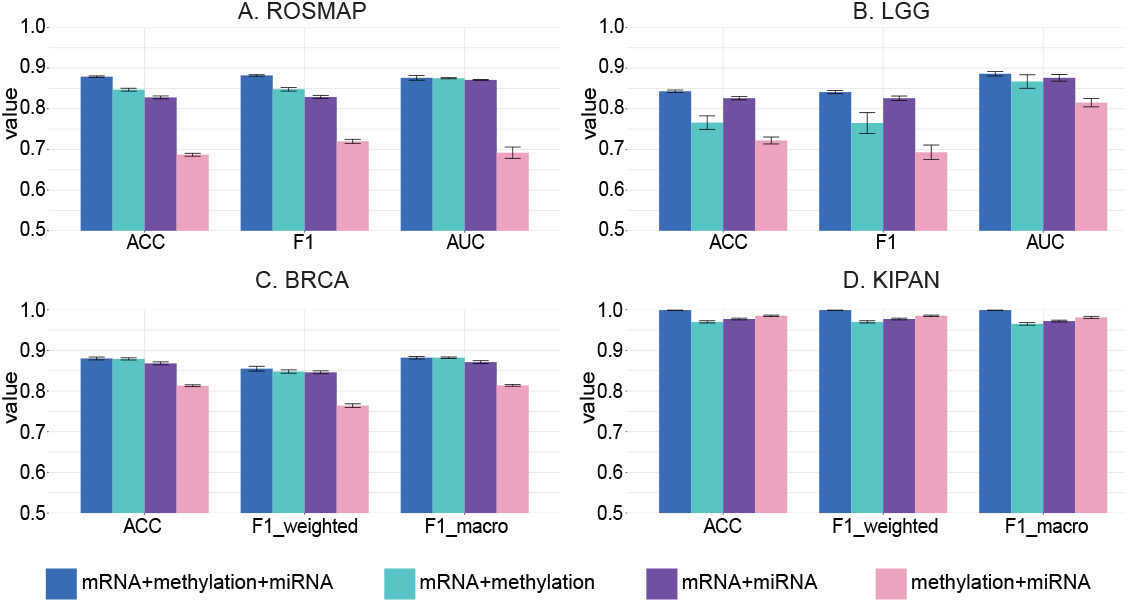
The efficacy of the proposed approach on different omics data combinations is assessed, presenting mean values and standard errors for comparison.

### 3.4. Robustness study involving comparisons with advanced methods

In the robustness experiments, we increase the masked ratio to heighten the uncertainty of a specified modality. This method assesses the robustness of our model by comparing the reduction in accuracy as the data becomes increasingly incomplete or uncertain. Figure 3 demonstrates that TEMINET exhibits superior stability, with consistently lower accuracy reduction ratios across all masked ratios, compared to Mogonet and Dynamic. The robustness of TEMINET is attributed to its use of a graph-based topology and an uncertainty-based trust mechanism. While Mogonet also employs a graph-based approach by constructing similarity graphs among samples and incorporating the VCDN trust mechanism, its performance is moderate. Conversely, the Dynamic method, which relies on an encoder network and a confidence-based trust mechanism, displays a greater decrease in accuracy, indicating reduced robustness relative to TEMINET. The findings underscore that TEMINET not only maintains lower accuracy reduction ratios but also exhibits stability across various levels of data masking, highlighting the effectiveness of its graph-based approach and uncertainty-based trust mechanism in preserving model robustness.

**Figure 3.**
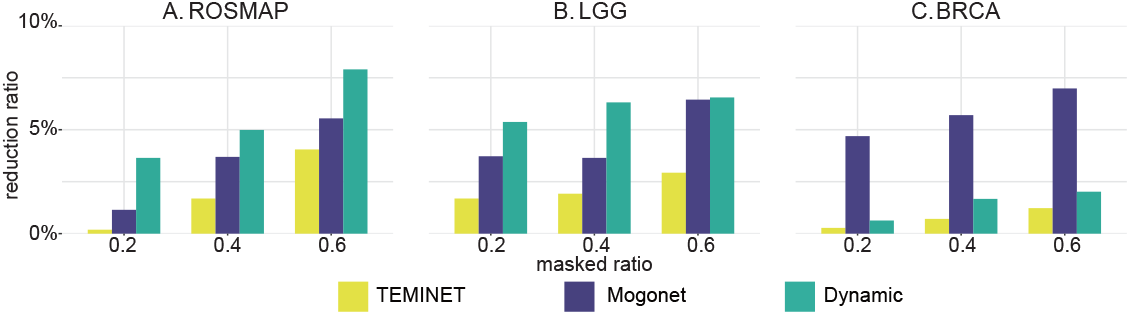
The robustness of the proposed approach is evaluated by comparing it with advanced methods. KIPAN is excluded from this comparison as it is relatively easy to classify.

### 3.5. Important biomarkers identified by TEMINET

In interpreting the model, the primary objective was to identify biomarkers of significance. As shown in Table 5, the five most discriminating biomarkers based on their differential values were reported. Those biomarkers with identical values received the same ranking. If the number of tied positions exceeded the threshold for reporting (i.e., five distinct ranks), a random set from the tied group was chosen to fulfill the report. Subsequently, we conducted a brief review of the existing literature to elucidate the biological significance and disease association of these identified biomarkers.

**Table 5.**
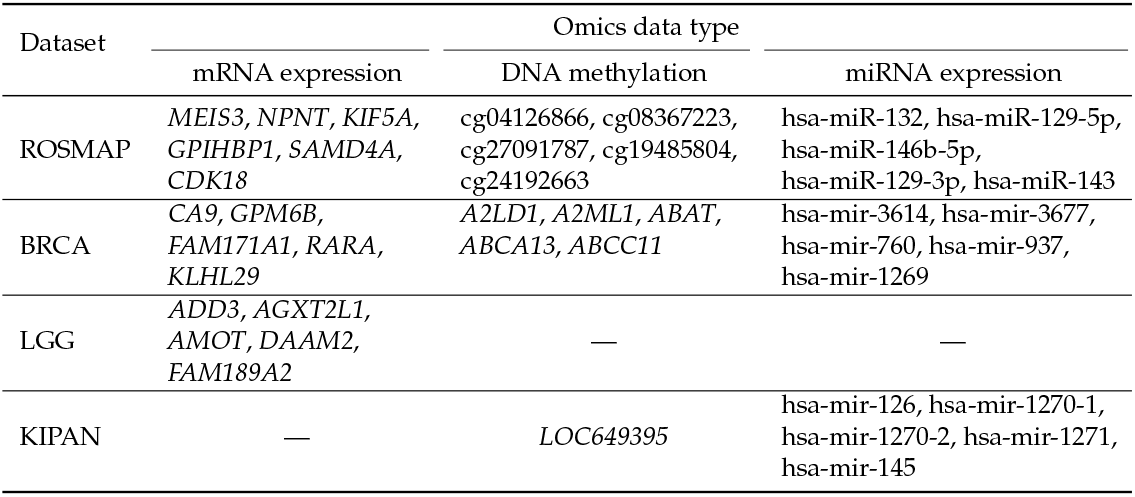
Top five significant disease-specific biomarkers identified by the TEMINET.

As shown in Table 5, existing advances in AD research have identified biomarkers associated with its pathogenesis. As the most significant mRNA biomarker of AD identified by the TEMINET, *MEIS3* was revealed by differential expression analysis to be considerably linked with cognitive decline and increased neurofibrillary tangle density [41,42]. Complementing this, cg19485804 (*NGEF*) emerged from LASSO regression analysis as another insight[43], notably associated with the APP mutation in mouse models. The downregulation of *NGEF* in the CVN-AD model suggested a critical role in modifying actin dynamics and consequently disrupting neuronal growth cone motility [44]. Furthermore, the microRNA miR-132 has been associated with the progression of A*β* and tau pathologies, with its reduced levels in circulation suggesting its utility as a potential diagnostic insight for AD. In the realm of breast cancer, *CA9* has been identified as a mRNA biomarker. A study has shown that BRCA patients with lower levels of *CA9* derive more benefit from adjuvant therapies, suggesting that *CA9* expression could be instrumental in tailoring patient-specific treatment plans [45]. The interaction between IGF2BP3, TRIM25, and miR-3614 delineated a novel regulatory pathway crucial for tumor cell proliferation. The protective role of IGF2BP3 in safeguarding TRIM25 mRNA from degradation and its influence on miR-3614 maturation presented new potential targets for therapeutic intervention in BRCA [46]. In LGG, our model did not reveal any methylation or miRNA biomarkers. However, *ADD3* was identified as the leading mRNA biomarker. Essential for actin cytoskeleton assembly, *ADD3* deficiency in GBM cells triggered pro-angiogenic signaling, enhancing VEGFR expression in endothelial cells, which could have implications for angiogenesis in LGG [47]. Suggested as a tumor suppressor and survival predictor on chromosome 10q, *ADD3* was valuable for prognostic assessments in LGG [48]. In kidney-related cancers, miR-126 has been recognized for its strong prognostic potential in clear cell renal cell carcinoma (ccRCC) [49], while miR-1271, markedly upregulated in ccRCC tissues, emerges as a significant marker for assessing disease severity [50]. These findings affirmed the robust capability of TEMINET in elucidating complex biological markers pertinent to disease mechanisms and therapy responsiveness.

## 4. Discussion and conclusion

The advancement in high-throughput techniques and individualized healthcare approaches has produced various supervised data collections critical for predictive applications such as pinpointing disease conditions, classifying tumor stages, and distinguishing cancer subtypes. The fusion of multi-omics information demonstrates enhanced efficacy in disease prediction compared to single omics approaches. For clinical applications, these integration models must not only provide precise diagnostic guidance but also cover a wide range of diseases. This underscores the necessity for models that exhibit both high accuracy and strong generalization across diverse medical conditions.

To achieve optimal multi-omics integration, the informativeness of modality representation has increasingly attracted attention. On the one hand, this informativeness reflects the quality of features specific to each omics type. This quality is not only contingent on the methodologies employed for feature representation but also on the inherent quality of the data. This is because data quality can be compromised during collection, storage, and processing, leading to potential loss and degradation. On the other hand, the informativeness of a modality significantly determines its contribution to the integration process. This contribution is measured by the extent to which data from a modality can complement or enhance the understanding that other omics types provide. It’s not merely about the quantity of data each modality brings but the unique biological insights it offers that cannot be captured by others. Therefore, evaluating the informativeness of each modality representation is essential for ensuring that the most informative and relevant features are utilized for better predictive accuracy and deeper biological understanding.

In this study, we introduced TEMINET, a framework optimized for the trustworthy integration of multi-omics datasets. The advanced performance of TEMINET was credited to its joint observation of intra-omics molecular interactions and inter-omics informativeness. The framework addressed multifaceted complexities in disease pathogenesis by amalgamating omics data with topological models. Our approach involves constructing individual graphs for each subject within each omics data. This data formation strategy allows us to leverage the inner topological information among intra-omics molecules obtained from disease-specific data to improve model performance. It emphasized the importance of the interplay between genetic factors in revealing the underlying causes of diseases. Evidence from investigations revealed consistent outperformance of TEMINET across multiple metrics in four distinct tasks. This confirms that enhancing intra-omics informativeness can significantly improve the application performance of trustworthy learning strategies in multi-omics integration. Moreover, the combination ablation experiment substantiated the hypothesis that the growing variety of omics data types had significantly improved integration capabilities. Applied across four diverse diseases, TEMINET enhanced the understanding of disease mechanisms and patient stratification, revealing biomarkers as potential insights and offering precise classifications that aided healthcare professionals in developing personalized therapeutic interventions based on these deeper insights into patient conditions.

However, the model exhibits several limitations. While it concentrates on specific omics interactions, the model might overlook the broader dynamics between different omics. This oversight could reduce AUC performance, indicating that a more balanced approach should be considered in future developments. The handling of complex omics-specific co-expression networks by the model, though effective in certain datasets, is computationally demanding. This is particularly evident in its performance with the BRCA dataset, indicating potential scalability issues for larger datasets. Moving forward, our efforts will focus on enhancing the algorithm of the model to integrate and interpret broad inter-omics patterns more effectively, aiming for a comprehensive understanding of complex disease mechanisms.

## Author Contributions

Conceptualization, Xiaohui Yao and Shan Cong; methodology, Haoran Luo and Hong Liang; investigation, Haoran Luo and Hongwei Liu; data preparation, Haoran Luo and Zhoujie Fan; writing—original draft, Haoran Luo and Shan Cong; writing—review and editing, Yanhui Wei, Xiaohui Yao, and Shan Cong. All authors have read and agreed to the published version of the manuscript.

## Data Availability Statement

The raw omics data of ROSMAP is available at https://doi.org/10.7303/syn2580853. The raw omics data of BRCA, LGG, and KIPAN are available at https://portal.gdc.cancer.gov/. Codes will be released soon.

## Disclaimer/Publisher’s Note

The statements, opinions and data contained in all publications are solely those of the individual author(s) and contributor(s).

